# Deep Coupled Kuramoto Oscillatory Neural Network (DcKONN): A Biologically Inspired Deep Neural Model for EEG Signal Analysis

**DOI:** 10.1101/2025.09.26.678831

**Authors:** Sayan Ghosh

**Affiliations:** Computer Science and Engineering, SRM AP, Amaravati 522240, Andhra Pradesh, India

**Keywords:** EEG, Classification, Kuramoto Oscillator, Deep Coupled Kuramoto Oscillatory Neural Network

## Abstract

Deep neural networks applied to signal processing tasks often need specialized architectural mechanisms to capture the temporal history of input signals. Traditional approaches include recurrent loops between layers, gated units, or tapped delay lines. However, biological brains exhibit much richer dynamics, characterized by activity across multiple frequency bands (alpha, beta, gamma, delta) and phenomena such as phase locking and synchronization. Standard Recurrent Neural Networks (RNNs) are limited in their ability to represent these complex dynamical features. In this work, we introduce a novel framework called the Deep Coupled Kuramoto Oscillatory Neural Network (DcKONN), which leverages networks of nonlinear Kuramoto oscillators trained in a deep learning paradigm. The DcKONN architecture has been applied to EEG signal classifier task. Simulation results demonstrate that the proposed oscillatory neural networks achieve superior or comparable classification accuracy compared to existing state-of-the-art models. Beyond performance improvements, these models also provide valuable neurobiological insights by naturally incorporating oscillatory dynamics into their architecture.

## I. INTRODUCTION

EEG records brain oscillations across delta, theta, alpha, beta, and gamma bands, each linked to functions like perception, memory, and cognition [1]. These rhythms facilitate resonant communication across neural populations, thereby supporting the transfer and integration of information.

Oscillatory Neural Networks (ONNs) employ nonlinear Hopf oscillators to merge biologically inspired rhythms with deep learning, offering efficient temporal modeling of EEG-like signals [2-6]. Beyond DONN based on Hopf oscillators, other oscillator-based frameworks have also been applied in EEG research—for instance, Kuramoto networks in the CAS regime can reproduce healthy and epileptic EEG with forecasts up to ∼5.8 seconds ahead at 9–11% error [7], while neural mass models like Jansen–Rit capture rhythms across all major frequency bands and complex dynamics such as photic driving [8].

In parallel, advances in computational power and deep learning have greatly enhanced EEG analysis, a field explored for decades using neural networks [9-11]. Overall, deep neural networks continue to drive progress in healthcare-oriented EEG applications.

## II. Methods

### A. Backgrounds

Hopf oscillators capture rich local amplitude–phase dynamics and power DONNs [2], but they lack explicit modeling of network-level phase interactions. Kuramoto networks, with coupling and Hebbian learning, directly represent synchronization and connectivity patterns critical for EEG analysis. They are computationally lighter, enabling scalable training while preserving biologically inspired mechanisms, such as phase-locking and modularity.

### B. Kuramoto Network Model

The mathematical description of a single Kuramoto neuron can be described by:

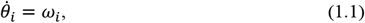

Where ω is the natural frequency of the oscillator. When an additive external signal drives a single Kuramoto oscillator, the state equation can be written: 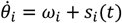.Where, *s*_*i*_(*t*) is the external driving force In general,

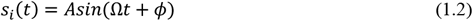

*A* is the amplitude of the externally applied signal, *Ω* is the angular frequency and *ϕ* phase. For our case it is an EEG signal.

The network of the Kuramoto Neuron model without an external driving force can be written:

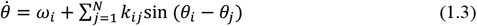

Where, *k*_*ij*_ is the real coupling coefficient, which encodes how j’th oscillator’s is influencing the ith oscillator. When a network of coupled Kuramoto oscillators is driven by an external signal, it is written by:

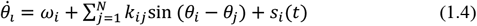

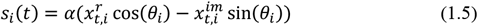

In a network, we have utilized ring topology with 2-neighbor coupling; the coupling term for oscillator *i* in the Kuramoto model is defined by its connections to the two immediate neighbours on either side. 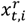 and 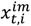 are the real and imaginary parts of the input projection (for our case, it is an EEG signal). *α* is the input scaler, which strengthens the driving force. *k*_*ij*_ is the learnable coupling coefficient whose weights have been updated using the biological Hebbian learning rule.

Now defined the adjacency matrix for *k*_*ij*_. If there are N oscillators are there, the adjacency matrix (*A*∈ ℝ^*N*×*N*^). We have used locally coupled ring topology in our study (Fig 1).

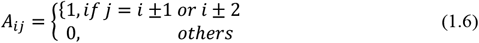

oscillatory network.

**Fig.1.**
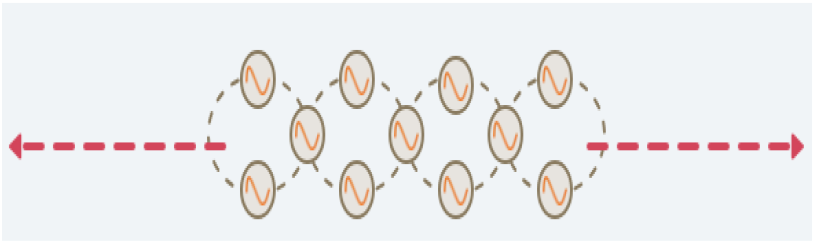
Graphical representation of a locally connected Kuramoto oscillatory network.

The effective coupling coefficient can be written:

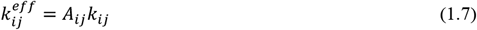

*k*_*ij*_ is the trainable weight,

Hebbian learning rule for the coupling coefficient

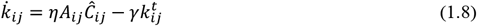

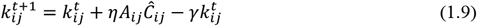

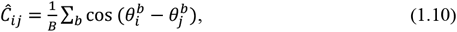

Where *η* is the Hebbian learning rate, *γ* acts as a decay constant. *B* is the batch size.

### C. Deep Coupled Kuramoto Oscillatory Neural Network (DcKONN)

Recently, Ghosh et al [2,4] proposed a deep oscillatory neural network based on Hopf oscillators, where in the oscillatory layers, oscillators are not coupled. In this model, we have proposed a coupled oscillatory network model (Fig. 2) where coupling coefficients are trained by a biological learning rule called Hebb’s rule.

**Fig.2.**
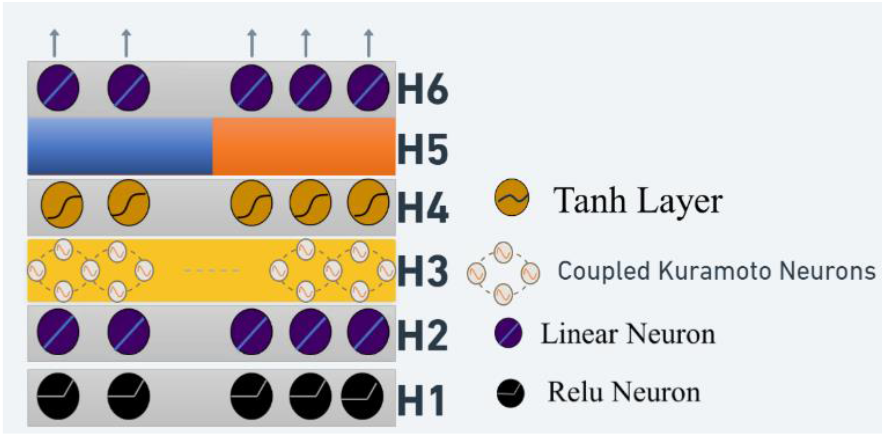
Model architecture.

Forward Propagation:

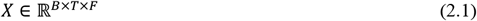

Where *X* is the EEG signal, B is the batch size, T is the time points, F is the number of channels.

At each time instant (dt) 1^st^ layer of the model gets input of *x*_*t*_

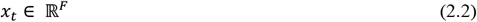

**1**^**st**^ **complex ReLU layer (H1)**: It has both real and imaginary pathways

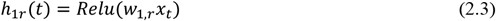

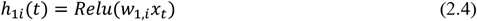

**2**^**nd**^ **complex dense layer (H2):**

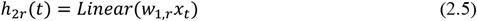

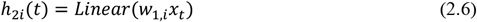

The real/imag inputs to the Kuramoto block:

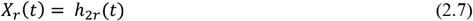

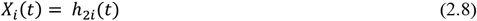

**Kuramoto Oscillatory Block (H3):**

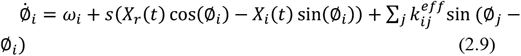

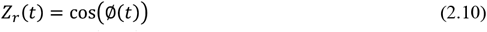

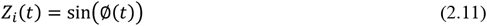

**Complex Tanh layer(H4):**

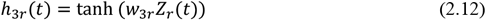

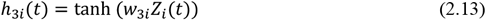

**Concatenation Layer(H5):**

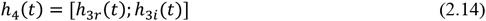

**Final Dense layer(H6):**

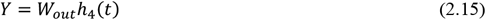

Also this deep oscillatory network is flexible; we can add or reduce the layers depending on our problems.

### D. Dataset and Pre-Processing

In this study, we used the EEGIS (Electroencephalogram Imagined Speech) dataset [13], selecting participants aged 20–30 years. EEG signals were recorded with a 14-channel Emotiv Epoc+ device (128 Hz) while participants, with eyes closed, imagined speaking one of eight Spanish words or remained in a relaxation state. Each trial lasted 8 seconds, segmented into overlapping 1-second windows (128 frames, 48-frame overlap), and filtered into five EEG bands: delta (0.1–4 Hz), theta (4–7 Hz), alpha (8–12 Hz), beta (12–30 Hz), and gamma (30–40 Hz). The dataset comprised 4,044 samples, with two tasks defined: binary classification of /Sí/ vs. /No/ (924 samples) and nine-class classification (all words plus relaxation). For both tasks, data were split into 80% training and 20% testing.

### E. Result: EEG classifier

The results section demonstrates that the Deep Coupled Kuramoto Oscillatory Neural Network achieves robust and high-performance classification of EEG signals. As shown in the training and validation curves (Fig 3), the model maintains low loss and high accuracy across 400 epochs, indicating stable convergence and minimal overfitting. Table 1 summarizes key architectural and training parameters, highlighting an efficient network configuration with biologically inspired components, such as Hebbian-learned coupling layers and a compact architecture (40 oscillators, 6,242 trainable parameters, 0.03 dropout, 0.001 learning rate, Adam optimizer). Table 2 presents a comprehensive performance comparison, showing that the model attains superior accuracy across all EEG frequency bands for both multi-class and binary (“yes vs. no”) tasks (e.g., theta: 98.86%; beta: 98.29%), outperforming classical approaches such as KNN with feature extraction. These findings confirm that integrating oscillator-based phase synchronization and Hebbian learning not only yields state-of-the-art accuracy but also offers interpretable neurobiological modeling for EEG classification tasks.

**Table 1.**
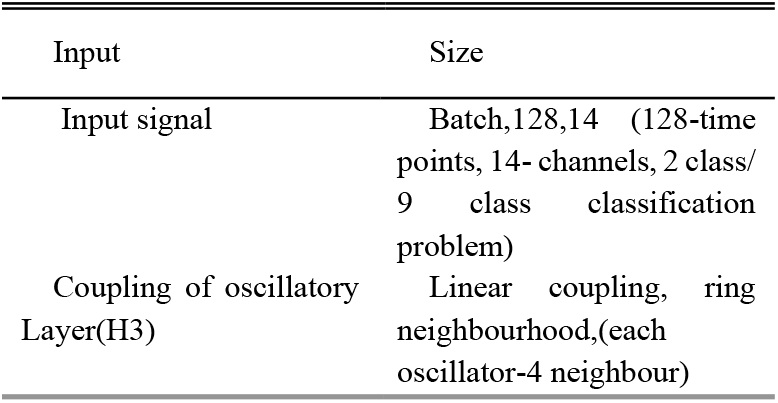
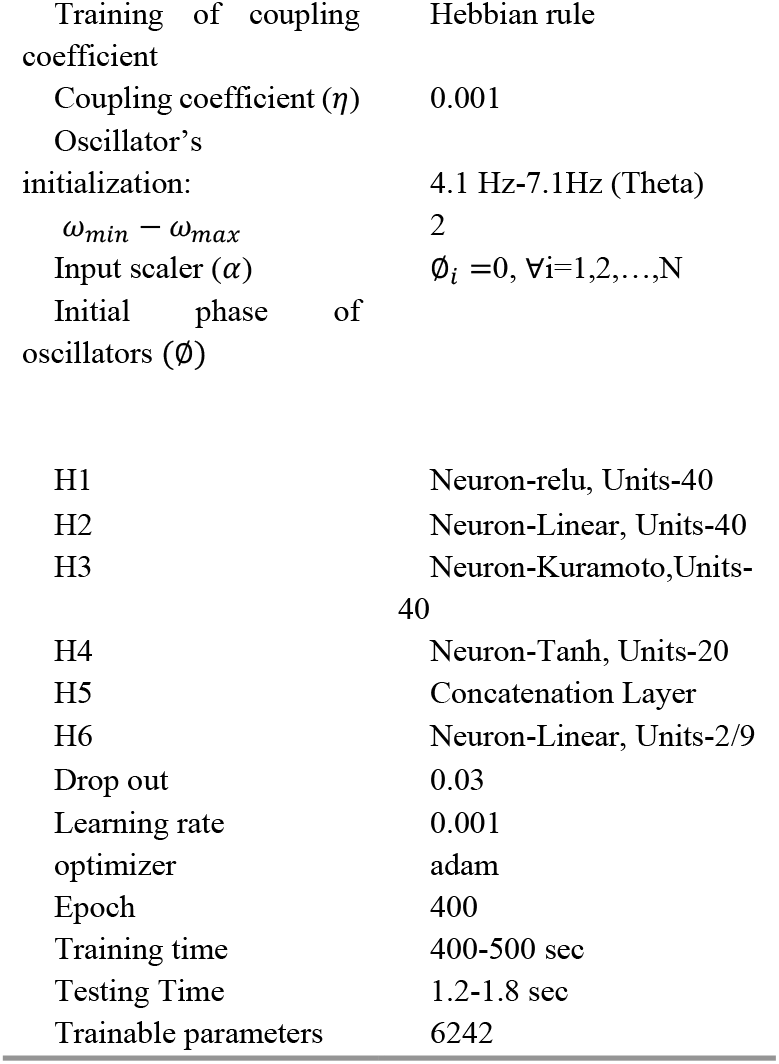
Network Parameters.

**Table 2.** Comparison Table.

| Symbol | Quantity <sup>a</sup> |
| --- | --- |
| Delta | All class: 80.78<br>Yes vs. no: 96.59 |
| Theta | All class: **87.98**<br>Yes vs. no: **98.86** |
| Alpha | All class: 76.38<br>Yes vs. no: 97.72 |
| Beta | All class: 79.12<br>Yes vs. no: **98.29** |
| gamma | All class: 51.2<br>Yes vs. no: 86.36 |
| KNN, feature extraction[13] | 96 |

**Fig.3.**
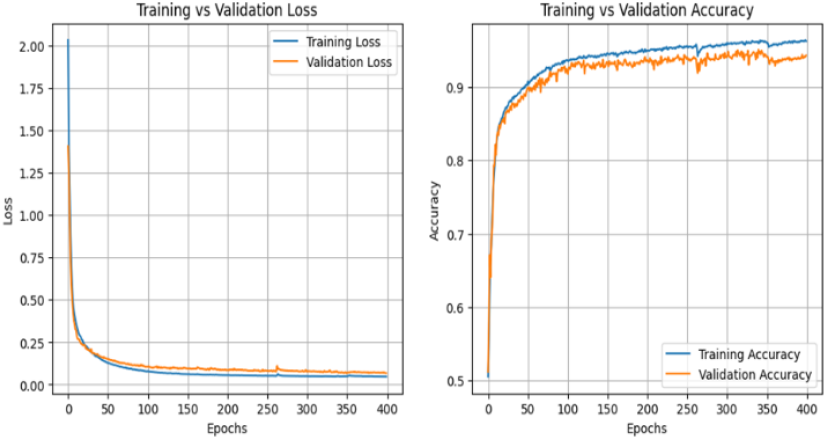
Network Performance.

Fig. 4 illustrates the learned coupling matrices from the Kuramoto– Hebbian layer for two EEG classes and their difference. Both Class 0 (left) and Class 1 (middle) exhibit strong local interactions along the diagonal due to the ring adjacency constraint, but their detailed coupling strengths differ. In Class 0, the strongest couplings are concentrated among early oscillators (0–10) and higher-index oscillators (28–38), whereas Class 1 shows more uniformly strong couplings across the diagonal, with notable reinforcement in the mid-range (10–25) and near oscillators 30–35. The difference matrix (right) highlights these distinctions, with red indicating stronger couplings in Class 1 and blue indicating stronger couplings in Class 0. Overall, Class 0 favors stronger early and high-index interactions, while Class 1 emphasizes more consistent and centrally focused couplings, revealing discriminative connectivity patterns learned by the model.

**Fig.4.**
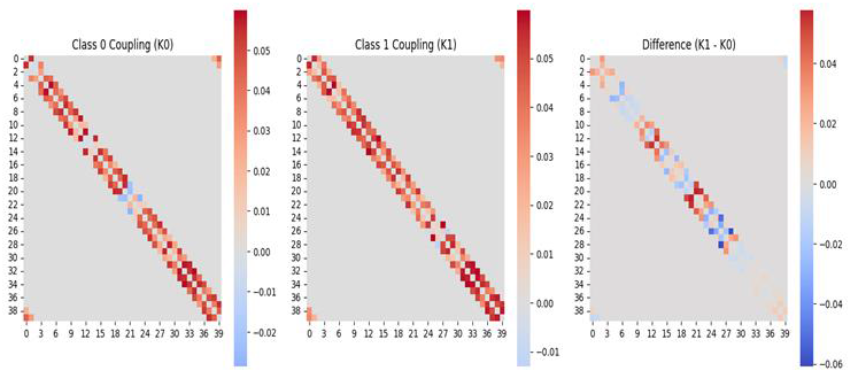
Learned coupling matrices from the Kuramoto–Hebbian layer for Class 0 and Class 1 EEG signals, and their difference matrix highlighting discriminative connectivity patterns.

The plot (Fig 5) displays the global synchrony *R*(*t*), where R(t) as a function of time for two classes: Class 0 (EEG) and Class 1 (Rest). The graph indicates a steady decline in synchrony for both classes, but Class 0 (EEG) retains a higher level of synchrony throughout the time period compared to Class 1 (Rest), which shows a more rapid decrease in synchrony. This suggests that the neural activity in Class 0, associated with EEG, exhibits more stable synchronization over time compared to the resting state of Class 1, where synchronization decreases more sharply. The divergence in synchrony patterns between the two classes highlights differences in brain dynamics under different conditions.

**Fig.5.**
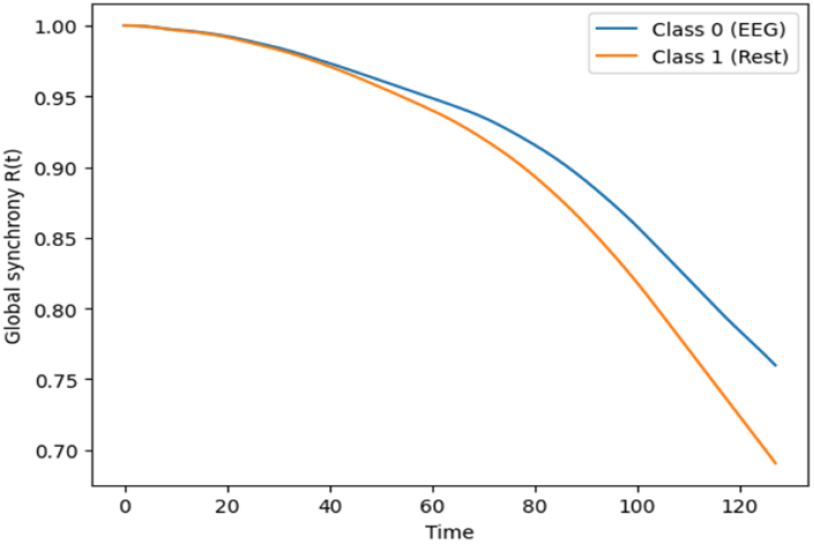
Temporal evolution of global synchrony (Kuramoto order parameter) and pairwise coordination (PLV) for Class 0 (EEG) and Class 1 (Rest) signals.

## III. CONCLUSION

The Deep Coupled Kuramoto Oscillatory Neural Network (DcKONN) represents a significant advance in biologically inspired EEG classification. By integrating Kuramoto oscillators with Hebbian coupling, the model not only delivers competitive accuracy across multiple EEG frequency bands, but also uncovers class-dependent synchrony patterns that offer interpretable neurobiological insights. This approach demonstrates that harnessing network-level phase synchronization and connectivity enables both robust signal classification and an improved understanding of underlying brain dynamics. The success of DcKONN highlights its potential for future application in clinical diagnostics and brain–computer interfaces, providing a powerful tool for both high-performance EEG analysis and the exploration of neurobiological mechanisms.

